# Rats Chasing the Dragon: A new heroin inhalation method

**DOI:** 10.1101/2023.08.09.552712

**Authors:** Arnold Gutierrez, Michael A. Taffe

**Affiliations:** Department of Psychiatry, University of California, San Diego; La Jolla, CA, USA

## Abstract

**Rationale:** Despite extensive human use of the inhalation route for ingesting opioids, models in rodents have mostly been limited to parenteral injection and oral dosing. Methods using electronic drug delivery systems (EDDS; “e-cigarettes”) have shown efficacy in rodent models but these do not faithfully mimic the most popular human inhalation method of heating heroin to the point of vaporization.

**Objective:** This study was designed to determine if direct volatilization of heroin hydrochloride delivers effective heroin doses to rodents.

**Methods:** Middle aged rats were exposed to vapor created by direct heating of heroin HCl powder in a ceramic e-cigarette type atomizer. Efficacy was determined with a warm water tail withdrawal nociception assay, rectal temperature and self-administration.

**Results:** Ten minutes of inhalation of vaporized heroin slowed response latency in a warm water tail withdrawal assay and increased rectal temperature in male rats, in a dose-dependent manner. Similar antinociceptive effects in female rats were attenuated by the opioid antagonist naloxone (1.0 mg/kg, s.c.). Female rats made operant responses for heroin vapor in 15-minute sessions, increased their response rate when the reinforcement ratio increased from FR1 to FR5, and further increased their responding when vapor delivery was omitted. Anti-nociceptive effects of self-administered volatilized heroin were of a similar magnitude as those produced by the 10-minute non-contingent exposure.

**Conclusions:** This study shows that “chasing the dragon” methods of inhalation of heroin can be modeled successfully in the rat. Inhalation techniques may be particularly useful for longer term studies deep into middle age of rat species.

## 1. Introduction

Public health crises involving non-medical use of opioids in Western countries date back at least to the 19th century, during which the inhalation of vapor created by heating opium (Kane, 1881; Kritikos, 1960) became associated with social and health problems, often with a racial blame component (Stephanie V. Ng, 2016). Although intravenous injection of heroin emerged through the 20^th^ century, other routes of administration, including the inhalation of vapors generated by heating heroin base or salts remain common. There is even some indication that inhalation use of heroin may come prior to a switch to injection for some users (Clatts et al., 2011; Cui et al., 2015; Rahimi-Movaghar et al., 2015; Syvertsen et al., 2017; Tyree et al., 2020), or remain the preferred method (Darke et al., 2004), perhaps due to perceptions of safety or cultural factors. The inhalation route might induce differences in the speed of brain entry, first pass metabolism, sequestration and release from non-brain tissues (Rook et al., 2006a; Rook et al., 2006b), any of which may alter the risks for developing a heroin use disorder, for example. Those who inhale heroin exhibited less severe psychiatric symptoms than those who inject heroin in one study (Wang and Liu, 2012) and inhalation is also associated with reduced risk of overdose (Darke et al., 2004). Inhalation may appear to be safer compared with intravenous injection, and indeed inhalation methods have been used in trials for a prescription modality of heroin assisted therapy for heroin dependence (Klous et al., 2005), encouraged in harm reduction attempts to decrease intravenous injection (Pizzey and Hunt, 2008; Stover and Schaffer, 2014) and investigated for delivering fentanyl for acute medical pain control (Vazda, Xia and Engqvist, 2021). Nevertheless, evidence shows that injection and inhalation have similarly increased severity of dependence compared with those who insufflate, particularly within the first 3 years of heroin use (Barrio et al., 2001).

Despite extensive human use of the inhalation route for ingesting opioids (Strang, Griffiths and Gossop, 1997), pre-clinical models in laboratory rodents have mostly been limited to parenteral injection, oral dosing or, for chronic exposure, the insertion of subcutaneous delivery systems (osmotic pumps, pellets, etc). One available prior study showed that the inhalation of volatilized drug produces analgesia / anti-nociceptive effects of heroin and other opioids in mice using a heated glass pipe approach and a tail-withdrawal assay (Lichtman, Meng and Martin, 1996). Interestingly morphine was 5-fold less potent than heroin when injected intravenously, but had similar potency when inhaled, further emphasizing a need for inhalation models. Additional studies show that monkeys will self-administer volatilized heroin (Mattox and Carroll, 1996; Mattox, Thompson and Carroll, 1997). These early reports did not result in subsequent broad use of inhalation techniques in other laboratories, potentially due to the difficulty of creating and maintaining inhalation apparatus suitable for laboratory use. This hurdle has been recently overcome by development and wide availability of e-cigarette style devices (aka Electronic Nicotine Delivery Systems or ENDS), which create a vapor from propylene glycol, vegetable glycerin and/or other constituents adulterated with nicotine. While these were originally marketed to deliver nicotine to human users, they were rapidly adapted for inhalation of cannabinoids, as well as numerous other psychoactive drugs, including opioids. These devices, better termed Electronic Drug Delivery Systems (EDDS), have been shown to be highly effective at delivering drugs via inhalation to laboratory rats and mice (Breit et al., 2022; Cooper, Akers and Henderson, 2021; Espinoza et al., 2022; Freels et al., 2020; Frie et al., 2020; Montanari et al., 2020; Moore et al., 2020; Nguyen et al., 2016a; Nguyen et al., 2016b; Smith et al., 2020), including opioids such as fentanyl, sufentanil, oxycodone and heroin (Gutierrez, Creehan and Taffe, 2021; Moussawi et al., 2020; Nguyen et al., 2019; Vendruscolo et al., 2018).

We have shown that heroin can be administered to rats via inhalation using a e-cigarette type device in which the heroin is dissolved in a propylene glycol vehicle, producing effects on nociception, locomotion and thermoregulation (Gutierrez, Creehan and Taffe, 2021), acting as a reinforcer of operant behavior in a self-administration paradigm (Gutierrez et al., 2020) and inducing lasting changes in anxiety-like behavior and nociception after repeated exposure during adolescence (Gutierrez et al., 2022a). While there are undoubtedly users who administer heroin in this manner (Blundell, Dargan and Wood, 2018; Breitbarth, Morgan and Jones, 2018), the inhalation of vapors generated by heating heroin directly (“chasing the dragon”) without an EDDS style liquid vehicle, is much more common. Thus, there is a need to create models by which the effects of inhaling directly volatilized heroin can be evaluated. Recent expansion of the EDDS market has produced devices which are designed for volatilizing cannabis preparations without a liquid vehicle. These devices are readily available online and in local vape shops and can therefore be readily adapted by any laboratory.

For this study, experiments were conducted to determine if the Honeystick XTRM 2.0 cartridge is capable of effectively delivering vapor from heroin HCl to rats. The Honeystick XTRM 2.0 cartridge is designed for cannabis wax, and it consists of a ceramic bowl fitted with two titanium heating elements. The *in vivo* assays selected assessed nociception (using the tail withdrawal assay), rectal temperature and self-administration. Preclinical rodent models rarely, if ever, model substance use in middle age adulthood following an extensive history of substances common in human opioid users such as nicotine or cannabis. Limited studies show that young rats self-administer more morphine than aged rats in the only available report (Bongiovanni et al., 2021). Another study reported the effect of morphine on brain reward thresholds was only altered at high doses in aged versus young adult rats (Jha, Knapp and Kornetsky, 2004). These studies were therefore conducted in middle aged rats with earlier life exposure to nicotine and/or Δ^9^-tetrahydrocannabinol (THC) as an improved translational model of middle age substance use.

## 2. Methods

### 2.1. Subjects

Male and female Sprague-Dawley rats were obtained from a commercial supplier (Envigo, Livermore, CA) and entered the laboratory at Post-Natal Day (PND) 23-25. Rats were housed in humidity- and temperature-controlled (23±2 °C) vivaria on 12:12 hour (reversed) light:dark cycles. Rats had *ad libitum* access to food and water in their home cages and all experiments were performed in the rats’ scotophase. The male (Gutierrez et al., 2022b) and female animals participated in the study after the conclusion of experiments previously described. All procedures were conducted under protocols approved by the Institutional Care and Use Committee of the University of California, San Diego.

#### Cohort 1

Groups (N=8 per group) of male Sprague-Dawley rats were exposed in twice daily 30-minute inhalation sessions with vapor from the Propylene Glycol (PG) vehicle or Nicotine (30 mg/mL in PG) from PND 31-40, and participated in additional studies from early adulthood onward, as described (Gutierrez et al., 2022b). In brief this involved evaluation of the locomotor effects of intermittent doses of nicotine (s.c.) from PND 74-105 in an open field and from PND 165-180 on an activity wheel. Animals were further assessed, starting PND 285, in nicotine (30mg/mL) EDDS vapor inhalation self-administration studies of 30 minute duration. Rats self-administered heroin (10, 50, 100 mg/mL) EDDS vapor in 14 sessions from PND 465-517. Experiments for this study were initiated on PND 536 for this cohort.

#### Cohort 2

Groups (N=8 per group) of female Sprague-Dawley rats were exposed in twice daily 30-minute inhalation sessions with vapor from the Propylene Glycol (PG) vehicle, Nicotine (60 mg/mL in PG), THC (100 mg/mL) or the Nicotine/THC combination. Rats were thereafter evaluated for temperature and antinociceptive responses to a THC injection (PND 80-81) and for locomotor effects of nicotine inhalation from ∼PND 114-128 on an activity wheel, in studies not currently described. Self-administration of nicotine EDDS vapor was conducted in 30 minute sessions every 2-4 days starting from PND 155-156. This included nicotine (30 mg/mL in the propylene glycol (PG) vehicle) from 22-26 and 30-33 weeks of age, nicotine (30 mg/ml) plus 5% menthol from weeks 34-35, nicotine (60 mg/mL) plus 5% menthol from weeks 36-40. The rats then received acute doses of heroin (0.0, 0.56, 1.0 mg/kg, s.c.) in a counter-balanced order, 3-4 days apart, prior to evaluation for rectal temperature and anti-nociception using the tail withdrawal assay from 42-43 weeks of age. Self-administration of e-cigarette vapor was re-initiated including heroin (50 mg/mL in the PG vehicle) from weeks 46-49, nicotine (30, 60 mg/mL in PG) and menthol (5% in PG) alone and in combination from weeks 50-53. For the present study rats (N=31) were evaluated from 55-57 weeks of age in the body temperature and nociception studies and a subset (N=6) thereafter in HCl vapor self-administration.

### 2.2 Apparatus

As previously described (Gutierrez et al., 2022b), during adolescent exposure nicotine and/or THC vapor was delivered twice a day in 30 minute sessions into sealed chambers (152 mm W X 178 mm H X 330 mm L; La Jolla Alcohol Research, Inc, La Jolla, CA, USA) through the use of e-vape controllers (Model SSV-3 or SVS-200; 58 watts, 0.24-0.26 ohms, 3.95-4.3 volts, ∼214 °F; La Jolla Alcohol Research, Inc, La Jolla, CA, USA) to trigger Smok Baby Beast Brother TFV8 sub-ohm tanks. Tanks were equipped with V8 X-Baby M2 0.25 ohm coils. MedPC IV software was used to schedule and trigger vapor delivery (Med Associates, St. Albans, VT USA). Sealed experimental chambers were ventilated with fresh air at a rate of ∼1-2 liters per minute via intake regulated by air meter and exhaust connected to laboratory house vacuum as in prior reports (Gutierrez et al., 2022c; Javadi-Paydar, Cole and Taffe, 2018; Nguyen et al., 2016a; Nguyen et al., 2016b).

For the present study, vapor was generated by a Honeystick extreme / XTRM 2.0 wax vape cartridge (Double Titanium Rod & Ceramic Donut Heater) triggered by a La Jolla Alcohol Research, Inc. SVS250 power device (or “mod”) set to 40 watts, with ∼2L/min air flow rate. Heroin HCl was placed in the bowl without any vehicle. Pilot studies determined that a 9 sec activation of the mod was required to sufficiently heat up the metal rod to produce drug volatilization (a “hit”). Studies were conducted under white light illumination to permit visual verification of the vapor cloud in the chambers. All visible drug was gone from the bowl after ∼4-6 of the 9 second power activation cycles, see below.

### 2.3 Drugs

Heroin (diamorphine HCl), nicotine and Δ^9^-tetrahydrocannabinol (THC) were administered by inhalation. Nicotine, THC and/or heroin were dissolved in propylene glycol (PG) for the adolescent vapor exposures, and for some self-administration. For this study, aliquots of 60 mg Heroin HCl were loaded in the tank for every session in which 1-4 hits were to be delivered to the chamber. This was increased to 80 mg for sessions in which 6 hits were to be delivered. Naloxone was dissolved in physiological saline and injected i.p…

### 2.4 Body Temperature and Nociception

Rats were evaluated for rectal temperature and tail withdrawal latency before, and then again after, the inhalation session. Sessions lasted 15 minutes during which heroin vapor was delivered at initiation and/or at 5 minutes. Air flow was ceased in the interval after each vapor delivery for 4:30 minutes. A 5-minute interval of airflow was conducted after 10 minutes to evacuate the chamber before animal removal for post-session testing.

Body temperature was determined by rectal measurement with a lubricated thermistor (VWR Traceable™ Digital Thermometer) as previously described (Gilpin et al., 2011). Nociception was assessed using a warm-water tail immersion assay, as described in (Javadi-Paydar et al., 2018; Nguyen et al., 2018a; Nguyen et al., 2018b). Briefly, the tail was inserted ∼3-4 cm into a warm (52°C) water bath and the latency for tail removal recorded with a stopwatch. A 15-second cutoff was used to avoid any potential for tissue damage.

Male rats (N=12; PND 536-552; ∼77-79 weeks of age) were examined for nociception and body temperature responses in three inhalation conditions. In our prior work with the EDDS system, dose has been manipulated by the concentration of drug in the PG vehicle and by total inhalation time. The latter approach slightly confounds time with vapor volume since hits were delivered every 5 minutes. For this initial study the inhalation time was held fixed (10 minutes) and the volume of vaporized heroin was varied. Rats were exposed to 1) Two 9-second hits at Time 0; 2) One 9-second hit at Time 0 and Time 5 min; or 3) Two 9 second hits at Time 0 and Time 5 min. Airflow was turned off after the hits, and turned back on 30 seconds prior to the Time 5 min hits to facilitate chamber re-filling. After a total of 10 minutes of exposure and 5 minutes of post-exposure chamber clearance, rats were removed and tested for anti-nociceptive and thermoregulatory responses. Rats were exposed to the inhalation conditions N=1 or N=2 per chamber, to facilitate the counterbalanced testing order.

Female rats (N=31; PND 387-401; ∼55-57 weeks of age) were examined for nociceptive and body temperature responses to inhalation of vaporized heroin HCl. For this study heroin exposure was varied by comparing two and three hits. The first subgroup (N=15) received two 9-second hits at Time 0 and Time 5 min of the session with 60 mg of heroin in the ceramic bowl. The second subgroup (N=16) received three 9-second hits at Time 0 and Time 5 min, with 80 mg of heroin HCl in the bowl to accommodate the extra hits. The two dose subgroups were made up of ∼equal numbers of animals from each of the original adolescent treatment groups. For this study the rats were injected with saline or naloxone (1.0 mg/kg, i.p.) prior to one of two inhalation sessions in a counterbalanced order. One rat in the second subgroup did not complete the two pre-treatment conditions thus N=15 in that group for analysis.

### 2.5 Self-administration

A subset of the female rats (N=6) were permitted to make nose-poke responses to obtain puffs of vaporized heroin HCl in 15-minute sessions. As noted, these rats had extensive prior experience responding for vapor deliveries of nicotine (30, 60 mg/mL in the PG vapor, 36 total sessions, 13 including 5% menthol) and seven sessions for heroin (50 mg/mL in the PG). There were three weeks of no self-administration sessions before starting the sessions in which Heroin HCl vapor puffs were delivered.

These studies were conducted at PND 410-429, or 58-60 weeks of age. Sessions were limited in duration at this stage because one load of the ceramic bowl only permitted up to 6 hits with high confidence of equivalent drug delivery. Vapor deliveries were initially available on a FR1 schedule for two sessions and a FR5 schedule thereafter. The devices were triggered for 9 seconds per reinforcer. A no-drug substitution session was inserted after the first four FR5 sessions and on the final two sessions, the reinforcer was vapor created from heroin dissolved in PG at a 50 mg/mL concentration, as described (Gutierrez et al., 2020). Tail withdrawal latency was determined before and after the second FR1 session and the no-drug session for comparison with the non-contingent data for this subset of animals from the above study.

### 2.6 Data Analysis

Data (body temperature, tail-withdrawal latency, vapor deliveries self-administered) were generally analyzed with two-way Analysis of Variance (ANOVA) including repeated measures factors for the Drug treatment condition and the Time after vapor initiation or injection. A mixed-effects model was used instead of ANOVA wherever there were missing datapoints. Any significant main effects were followed with post-hoc analysis using Tukey (multi-level factors) or Sidak (two-level factors) procedures. The criterion for inferring a significant difference was set to p<0.05. All analysis used Prism 8 or 9 for Windows (10.0.0; GraphPad Software, Inc, San Diego CA).

## 3. Results

### 3.1 Body Temperature and Nociception

Inhalation of heroin slowed tail withdrawal in a dose dependent manner (**Figure 1A**) and produced an increase in rectal temperature (**Figure 1B**) in male rats. The statistical analysis confirmed significant effects of the factors of Time (before/after inhalation) (F (1, 11) = 173.0; P<0.0001), of vapor condition (F (2, 22) = 11.87; P<0.0005) and of the interaction (F (2, 22) = 11.92; P<0.0005) for tail withdrawal latency (**Figure 1A**). The Tukey post-hoc test confirmed that post-exposure withdrawal latency was longer for the double hit at times 0 and 5 minutes, compared with both other conditions. There was only a significant effect of the Time (F (1, 11) = 23.43; P<0.001) on rectal temperature (**Figure 1B**), and the post-hoc test confirmed latencies were slower post-inhalation for each test condition. There was no significant effect of the original adolescent treatment group (PG: N=5; Nic30: N=7) on post-session tail withdrawal or rectal temperature following heroin inhalation confirmed in a follow-up analysis which included a factor for adolescent treatment group.

**Figure 1:**
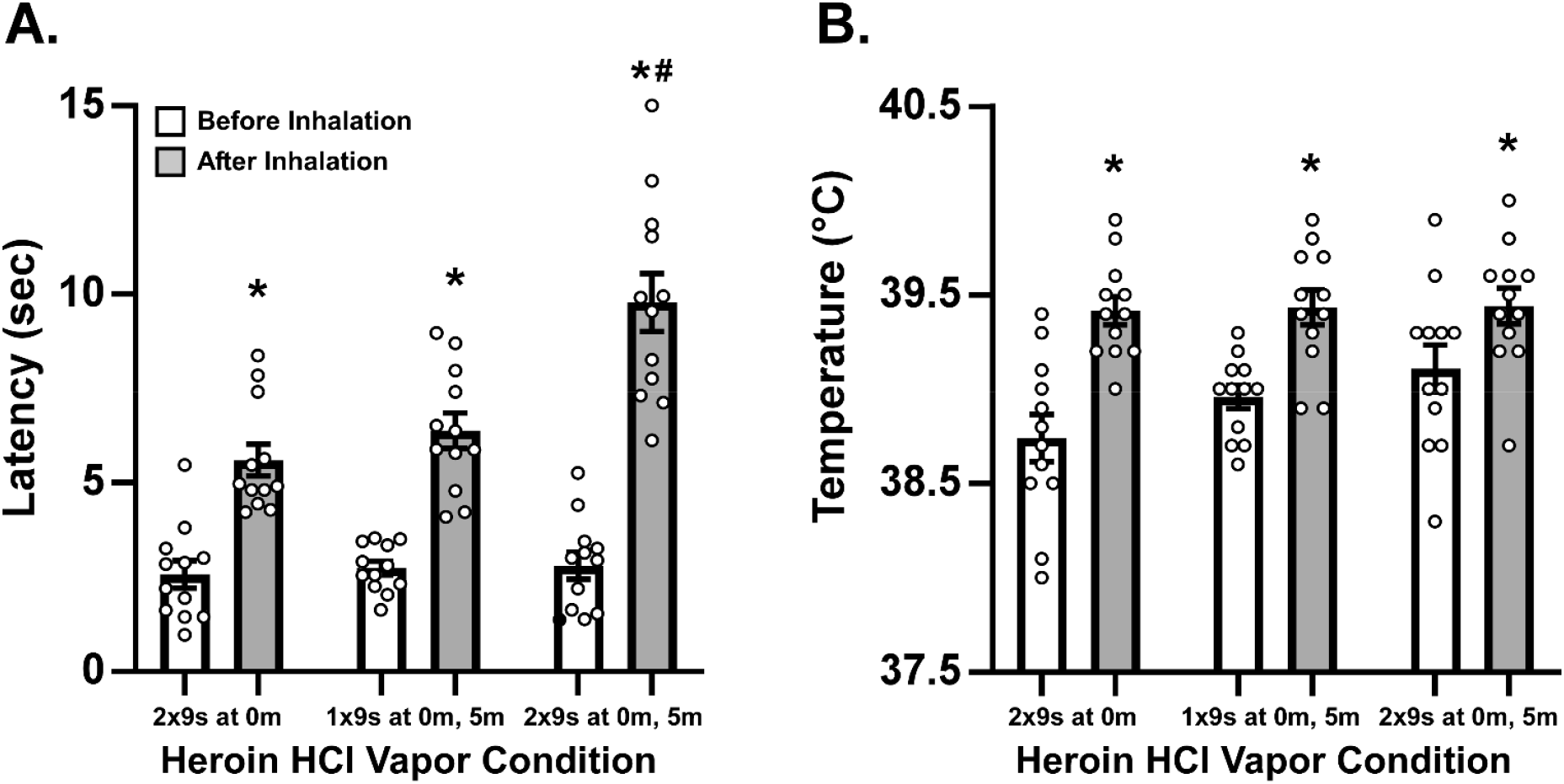
Mean (N=12; ±SEM) A) tail withdrawal latency and B) rectal temperature for male rats exposed to 10-minute inhalation sessions. A significant difference from the pre-session measurement is indicated with * and a difference from both other post-session measurements with #.

### 3.1 Antagonist Challenge of Heroin HCl Vapor Effects

Heroin inhalation slowed tail withdrawal latency and decreased the rectal temperature of female rats (**Figure 2**). The naloxone pre-treatment partially attenuated the anti-nociceptive effect but not the change in body temperature. The initial analysis included factors for dosing subgroup, for pre-inhalation Treatment (Saline vs Naloxone) and for Time (before and after inhalation). This analysis confirmed significant effects of Time [F (1, 28) = 132.0; P<0.0001], of Treatment Condition [F (1, 28) = 11.79; P<0.005] and the interaction of Time with Treatment Condition [F (1, 28) = 12.01; P<0.005], but no impact of HCl dose alone or in interaction with other factors for tail withdrawal latency. The analysis of body temperature confirmed only a significant effect of Time [F (1, 28) = 13.90; P<0.001]. Thus, the addition of an extra 9-sec hit at time 0 and time 5 minutes of the session did not functionally alter the delivered heroin dose.

**Figure 2:**
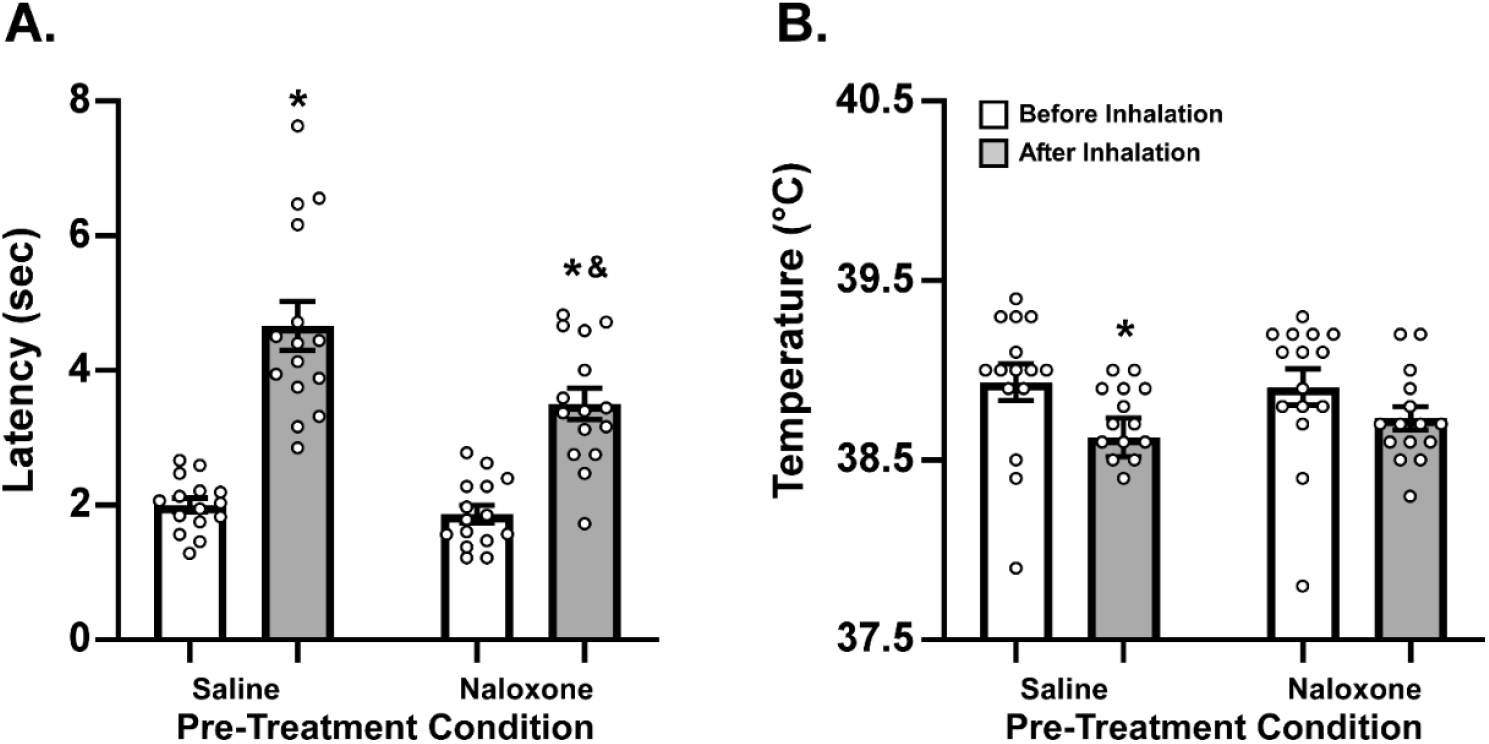
Mean (N=30; ±SEM) A) tail-withdrawal latency and B) rectal temperature of female rats before and after inhalation of heroin, following injection with Saline or Naloxone (1.0 mg/kg, s.c.). A significant difference between Before and After inhalation is indicated by * and a significant difference between Saline and Naloxone conditions with &.

The two factor ANOVA, collapsed across HCl dosing subgroup, confirmed that there was a significant effect of measurement Time [F (1, 14) = 83.23; P<0.0001], of Treatment Condition [F (1, 14) = 11.22; P<0.005] and the interaction [F (1, 14) = 10.87; P<0.01] on tail withdrawal latency. The Tukey post-hoc test confirmed there were significant differences between pre and post inhalation measurements for both Saline and Naloxone conditions, as well as between the two post-inhalation measurements. Analysis of the rectal temperatures confirmed that there was a significant effect of measurement Time [F (1, 14) = 83.23; P<0.0001], but not of Treatment Condition or of the interaction. There was no significant effect of the original adolescent treatment group (N=8, except PG group N=6) alone or in interaction with other factors on post-session tail withdrawal or rectal temperature following heroin inhalation confirmed in a follow-up analysis, which included a factor for adolescent treatment group.

### 3.2 Self-administration of Heroin HCl Vapor

A subset (N=6) of the female rats were permitted to respond (FR1) for heroin HCl vaporized or in a session in which the vaporizer was disconnected (No-tank condition). The rats responded voluntarily for vaporized heroin HCl, and engaged in drug-seeking behavior when the response contingency was increased to FR5 or when no vapor was delivered (**Figure 3**). Changing the reinforcer to puffs of PG adulterated with heroin (50 mg/mL) using the EDDS method resulted in similar numbers of reinforcers acquired, and the group maintained a mean of over 80% of responses directed at the drug-associated manipulandum across the study. Analysis with one way ANOVAs confirmed significant differences in reinforcers acquired [F (3.391, 16.96) = 7.02; P<0.005], correct responses [F (2.572, 12.86) = 7.87; P<0.005], and correct responses during the timeout interval [F (1.368, 6.842) = 9.67; P<0.05], but not in percent correct responses [F (3.335, 16.68) = 1.62; P=0.2196], across sessions. The post-hoc analysis further confirmed that introduction of the FR5 contingency increased responding on the drug-associated manipulandum and decreased reinforcers acquired. When vapor delivery was omitted, this increased both drug-associated responding and the number of reinforcers acquired.

**Figure 3:**
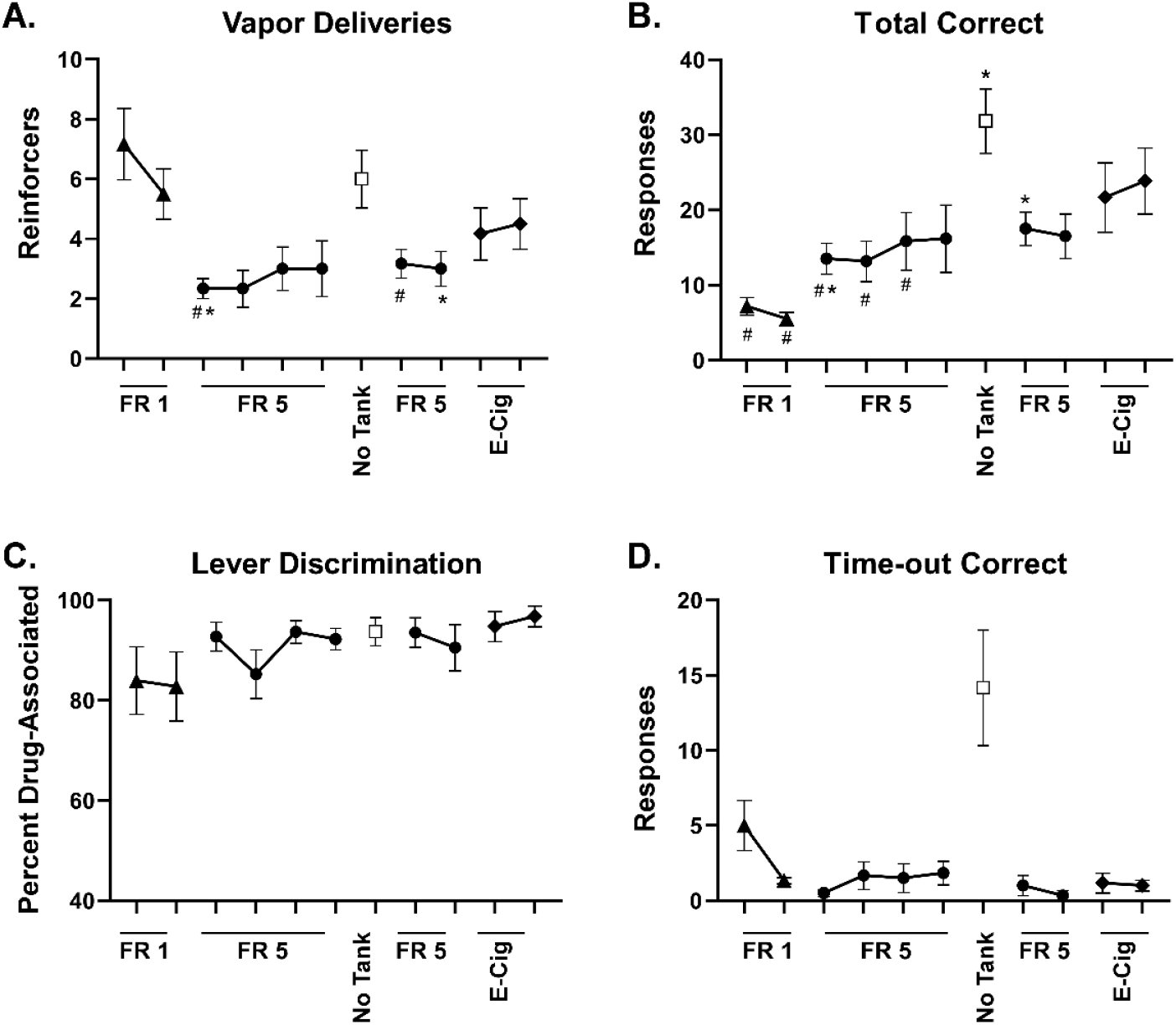
Mean (N=6; ±SEM) A) reinforcers acquired, B) correct responses, C) percent correct responses and D correct responses during the timeout interval for female rats. A significant difference from the first session is indicated with * and a significant difference from the no-tank condition is indicated with #.

**Figure 4:**
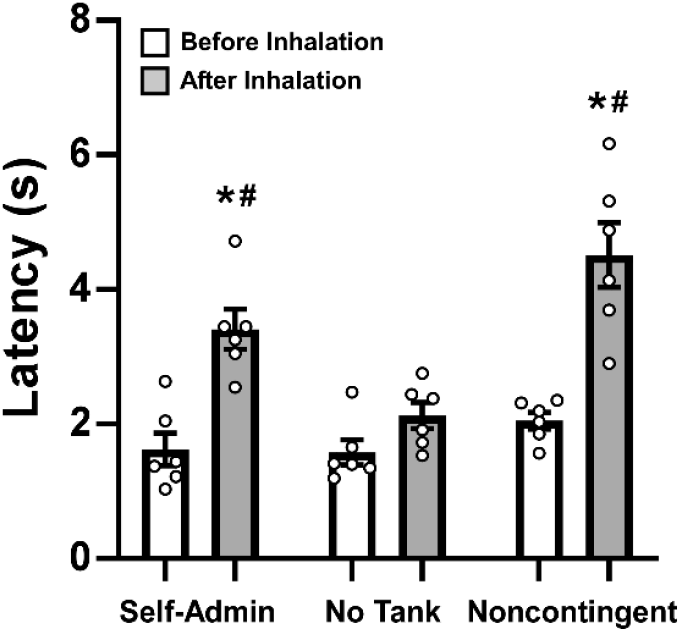
Mean (N=6; ±SEM) tail withdrawal latency before and after self-administration sessions with the heroin HCl or with the tank detached is compared with latencies for the same individuals from the noncontingent exposure in Experiment 2. A significant difference from before inhalation is indicated with * and a significant difference from the no-tank condition is indicated with #.

#### Antinociceptive effects of Heroin Inhalation Self-Administration

Tail withdrawal latencies observed before after the second self-administration session and the no-tank session were contrasted with their tail flick in the above non-contingent heroin HCl experiment. The ANOVA confirmed there was a significant effect of Time [F (1, 5) = 25.36; P<0.005] and of Test Condition (self-administration v no-tank self-administration v non-contingent) [F (2, 10) = 25.74; P=0.0001] and of the interaction of factors [F (2, 10) = 8.10; P<0.01]. The Tukey post-hoc test of all possible comparisons confirmed significant differences between the pre/post in both active drug conditions but not the No-tank self-administration condition. Likewise, a significant difference between the post-session latencies after the No-tank self-administration and each active drug condition was confirmed. There was no significant difference in post-session tail-withdrawal latency after heroin HCl self-administration compared with the non-contingent heroin HCl study confirmed.

## 4. Discussion

The present data show that direct volatilization of heroin HCl using a ceramic bowl e-cigarette device is capable of delivering a physiologically significant dose to rats via inhalation. Inhalation exposure sessions of 10 minutes duration produced anti-nociceptive effects in both male and female rats, an increase in male rat body temperature and a decrease in female rat body temperature. This pattern is similar to effects of inhalation of heroin using a traditional e-cigarette vehicle (PG) approach that were previously reported (Gutierrez, Creehan and Taffe, 2021; Gutierrez et al., 2022c). Furthermore, delivery of directly volatilized heroin reinforced operant responding in a self-administration paradigm. Thus, the human heroin use pattern termed “chasing the dragon” can be effectively modeled in laboratory rats.

The increase in rectal temperature that was observed in the male rats is similar to an effect observed in male and female rats after 15 minutes of Electronic Drug Delivery System (EDDS) heroin vapor inhalation (Gutierrez, Creehan and Taffe, 2021), and is consistent with reports that skin temperature is elevated in experienced human users during controlled tests of inhaled heroin (Hendriks et al., 2001; Rook et al., 2006b). Female rats’ temperature decreased after 30 minutes of EDDS heroin exposure (Gutierrez, Creehan and Taffe, 2021), thus the decrease in female rats’ temperature observed in this study may reflect delivery of a higher dose per unit inhalation time. It is also the case that temperature responses in rats can be biphasic, as an initial decrease 30-60 minutes after EDDS vapor inhalation was followed by a modest increase in temperature 90-120 minutes after inhalation (Gutierrez, Creehan and Taffe, 2021). The timing of this may depend on dose, animal age or other unknown factors.

The magnitude of effect on tail withdrawal latency in the male rats was similar to that produced by the inhalation of heroin (50 mg/mL concentration, 30 min) via the EDDS approach (Gutierrez, Creehan and Taffe, 2021; Gutierrez et al., 2022c). The comparatively smaller magnitude in the female group (**Figure 2**) is somewhat discordant with prior results, since female rats appeared to exhibit greater anti-nociceptive effects of heroin inhalation by EDDS (Gutierrez, Creehan and Taffe, 2021; Gutierrez et al., 2022c). It is possible that the middle age of these rats is associated with differences in antinociceptive response and thus future targeted studies may be warranted.

Self-administration by the inhalation method of drug delivery is a relatively recent approach, however EDDS self-administration has been described for several drugs including heroin, fentanyl and sufentanil (Gutierrez et al., 2020; McConnell et al., 2021; Moussawi et al., 2020; Vendruscolo et al., 2017). The present study was limited and conducted with animals that had an extensive history of self-administration of nicotine vapor and some experience with heroin vapor substitution. Here, it was demonstrated that animals maintained responding for directly volatilized heroin vapor, exhibited mean response ratios of over 80% on the drug-associated manipulandum, increased response rates when the schedule of reinforcement was increased from FR1 to FR5 and further increased their responding, including during the time-out interval, when vapor was no longer delivered. Furthermore, the self-administration of heroin to a subjective satiety level produced anti-nociceptive effects of a similar magnitude as were produced with involuntary dosing. Overall, this pattern of results is consistent with the volatilized heroin serving as a reinforcer within the expectations of a self-administration paradigm.

There are few reports of opioid effects in middle-age or aged rats. One study shows decreased intravenous self-administration of morphine in 20-24 month old Wistar rats compared with young adult rats (Bongiovanni et al., 2021); this is a range close to the lifespan of this strain. Morgan and colleagues reported that aged male F344/BNF_1_ rats are less sensitive than 12 month old F344/BNF_1_ rats to the antinociceptive effects of morphine in a 45°C floor preference assay (Morgan et al., 2012), and Jha and colleagues (2004) reported that aged rats are less likely to exhibit decreased reward thresholds in an intracranial self-stimulation reward paradigm after high doses of morphine, compared with younger adult rats. These latter studies (Jha, Knapp and Kornetsky, 2004; Morgan et al., 2012), however, used a F344/BNF1 strain hybrid rat that is more robust than most strains at ∼20-24 months and can live to 36 months (Agbas, Zaidi and Michaelis, 2005). This questions whether it is merely an experimentally tractable model of aged rats (many strains are nearing the end of life after 24 months and have unstable health) or a model of middle-aged rats which should be viewed as more similar to the 12-15 month interval in other strains. In the present study, in Cohort 1, one male was euthanized at PND 473 and one at PND 490 for development of abcess or tumor; One female from that study (not used here) was euthanized at PND 364 for a solid mass, all other male and female rats survived in appropriate health condition to at least PND 511. In Cohort 2, one female was euthanized at PND 319 for a solid mass, one died overnight at PND 398 of unknown causes. All other animals survived to at least PND 430. In short, the focus here was on rats in middle age, and not on late stage aging. Although this was a convenience sample of middle-aged rats that had been used in studies focused on the lasting effects of adolescent inhalation of nicotine, THC and/or the combination via a EDDS vapor inhalation approach, there is translational validity for middle aged human populations. A history of exposure to nicotine and cannabis is common in human opioid users.

One advantage of the described approach is that it is readily adapted by any interested laboratory since the ceramic bowl type of e-cigarette canister is now broadly commercially available. As has been the case with the EDDS vapor method, it is relatively simple for interested laboratories to set up sealed rat chambers with appropriate vacuum-controlled ventilation. This study outlines some initial parameters for drug delivery but, much as with EDDS methods, the parameters can be modified to fit various experimental needs. As a more general point, and particularly in terms of the self-administration experiment, it is unlikely that subject viability (and even survival) would be this high with intravenous catheterization approaches. Catheters become non-patent, ports can be chewed off by cagemates and/or subjects can experience external port rejection and other health complications. Vapor inhalation methods can avoid such health complications, as well as any difficulties that may occur in the vascular catheterization.

In conclusion, this study provides initial validation for a rat model of chasing the dragon. Inhalation of the vapor created by direct heat volatilization of heroin HCl altered nociception, body temperature and reinforced operant responding.

## Acknowledgements

The authors are grateful to Keven Creehan for expert technical assistance with the studies. The authors are also grateful to Maury Cole of La Jolla Alcohol Research, Inc. for provision of equipment necessary for volatilizing the heroin HCl. The study was conducted with the support of a UCSD Chancellor’s Post-doctoral Fellowship (AG), UCSD IRACDA (K12 GM068524; AG), and the Tobacco Related-Disease Research Program (TRDRP; T31IP1832). The TRDRP had no role in study design, collection, analysis and interpretation of data, in the writing of the report, or in the decision to submit the paper for publication. The authors declare no additional financial conflicts which affected the conduct of this work.

## Notes

### Competing Interest Statement

The authors are grateful to La Jolla Alcohol Research, Inc. for provision of equipment necessary for volatilizing the heroin HCl.

## Literature Cited

Agbas, A., Zaidi, A., Michaelis, E.K., 2005. Decreased activity and increased aggregation of brain calcineurin during aging. Brain Res 1059(1), 59–71 doi: 10.1016/j.brainres.2005.08.008.

Barrio, G., De La Fuente, L., Lew, C., Royuela, L., Bravo, M.J., Torrens, M., 2001. Differences in severity of heroin dependence by route of administration: the importance of length of heroin use. Drug and alcohol dependence 63(2), 169–177 doi: Doi 10.1016/S0376-8716(00)00204-0.

Blundell, M., Dargan, P., Wood, D., 2018. A cloud on the horizon-a survey into the use of electronic vaping devices for recreational drug and new psychoactive substance (NPS) administration. QJM 111(1), 9–14 doi: 10.1093/qjmed/hcx178.

Bongiovanni, A.R., Peer, K., Carpenter, R.E., Ellis, A.S., Duggan, M.R., Parikh, V., Wimmer, M.E., 2021. Aging reduces the sensitivity to the reinforcing efficacy of morphine. Neurobiol Aging 97, 28–32 doi: 10.1016/j.neurobiolaging.2020.09.020.

Breit, K.R., Rodriguez, C.G., Hussain, S., Thomas, K.J., Zeigler, M., Gerasimidis, I., Thomas, J.D., 2022. A Model of Combined Exposure to Nicotine and Tetrahydrocannabinol via Electronic Cigarettes in Pregnant Rats. Front Neurosci 16, 866722 doi: 10.3389/fnins.2022.866722.

Breitbarth, A.K., Morgan, J., Jones, A.L., 2018. E-cigarettes-An unintended illicit drug delivery system. Drug Alcohol Depend 192, 98–111 doi: 10.1016/j.drugalcdep.2018.07.031.

Cooper, S.Y., Akers, A.T., Henderson, B.J., 2021. Flavors Enhance Nicotine Vapor Self-administration in Male Mice. Nicotine Tob Res 23(3), 566–572 doi: 10.1093/ntr/ntaa165.

Darke, S., Hetherington, K., Ross, J., Lynskey, M., Teesson, M., 2004. Non-injecting routes of administration among entrants to three treatment modalities for heroin dependence. Drug Alcohol Rev 23(2), 177–183 doi: 10.1080/095952304100017044163.

Espinoza, V.E., Giner, P., Liano, I., Mendez, I.A., O’Dell, L.E., 2022. Sex and age differences in approach behavior toward a port that delivers nicotine vapor. J Exp Anal Behav 117(3), 532–542 doi: 10.1002/jeab.756.

Freels, T.G., Baxter-Potter, L.N., Lugo, J.M., Glodosky, N.C., Wright, H.R., Baglot, S.L., Petrie, G.N., Yu, Z., Clowers, B.H., Cuttler, C., Fuchs, R.A., Hill, M.N., McLaughlin, R.J., 2020. Vaporized Cannabis Extracts Have Reinforcing Properties and Support Conditioned Drug-Seeking Behavior in Rats. J Neurosci 40(9), 1897–1908 doi: 10.1523/JNEUROSCI.2416-19.2020.

Frie, J.A., Underhill, J., Zhao, B., de Guglielmo, G., Tyndale, R.F., Khokhar, J.Y., 2020. OpenVape: An Open-Source E-Cigarette Vapor Exposure Device for Rodents. eNeuro 7(5) doi: 10.1523/ENEURO.0279-20.2020.

Gilpin, N.W., Wright, M.J., Jr., Dickinson, G., Vandewater, S.A., Price, J.U., Taffe, M.A., 2011. Influences of activity wheel access on the body temperature response to MDMA and methamphetamine. Pharmacol Biochem Behav 99(3), 295–300 doi: S0091-3057(11)00139-0 [pii] 10.1016/j.pbb.2011.05.006.

Gutierrez, A., Creehan, K.M., Taffe, M.A., 2021. A vapor exposure method for delivering heroin alters nociception, body temperature and spontaneous activity in female and male rats. Journal of neuroscience methods 348, 108993 doi: 10.1016/j.jneumeth.2020.108993.

Gutierrez, A., Harvey, E.L., Creehan, K.M., Taffe, M.A., 2022a. The long-term effects of repeated heroin vapor inhalation during adolescence on measures of nociception and anxiety-like behavior in adult Wistar rats. Psychopharmacology (Berl) 239(12), 3939–3952 doi: 10.1007/s00213-022-06267-6.

Gutierrez, A., Nguyen, J.D., Creehan, K.M., Grant, Y., Taffe, M.A., 2022b. Adult consequences of repeated nicotine vapor inhalation in adolescent rats. bioRxiv, 2022.2011.2017.516984 doi: 10.1101/2022.11.17.516984.

Gutierrez, A., Nguyen, J.D., Creehan, K.M., Javadi-Paydar, M., Grant, Y., Taffe, M.A., 2022c. Effects of combined THC and heroin vapor inhalation in rats. Psychopharmacology (Berl) 239(5), 1321–1335 doi: 10.1007/s00213-021-05904-w.

Gutierrez, A., Nguyen, J.D., Creehan, K.M., Taffe, M.A., 2020. Female rats self-administer heroin by vapor inhalation. Pharmacol Biochem Behav 199, 173061 doi: 10.1016/j.pbb.2020.173061.

Hendriks, V.M., van den Brink, W., Blanken, P., Bosman, I.J., van Ree, J.M., 2001. Heroin self-administration by means of ‘chasing the dragon’: pharmacodynamics and bioavailability of inhaled heroin. Eur Neuropsychopharmacol 11(3), 241–252 doi: 10.1016/s0924-977x(01)00091-8.

Javadi-Paydar, M., Cole, M., Taffe, M.A., 2018. Effects of Nicotine and THC Vapor Inhalation Administered by An Electronic Nicotine Delivery System (ENDS) in Male Rats. bioRxiv, 418699 doi: 10.1101/418699.

Javadi-Paydar, M., Nguyen, J.D., Kerr, T.M., Grant, Y., Vandewater, S.A., Cole, M., Taffe, M.A., 2018. Effects of Delta9-THC and cannabidiol vapor inhalation in male and female rats. Psychopharmacology (Berl) 235(9), 2541–2557 doi: 10.1007/s00213-018-4946-0.

Jha, S.H., Knapp, C.M., Kornetsky, C., 2004. Effects of morphine on brain-stimulation reward thresholds in young and aged rats. Pharmacol Biochem Behav 79(3), 483–490 doi: 10.1016/j.pbb.2004.08.014.

Klous, M.G., Huitema, A.D., Rook, E.J., Hillebrand, M.J., Hendriks, V.M., Van den Brink, W., Beijnen, J.H., Van Ree, J.M., 2005. Pharmacokinetic comparison of two methods of heroin smoking: ‘chasing the dragon’ versus the use of a heating device. Eur Neuropsychopharmacol 15(3), 263–269 doi: 10.1016/j.euroneuro.2004.12.001.

Lichtman, A.H., Meng, Y., Martin, B.R., 1996. Inhalation exposure to volatilized opioids produces antinociception in mice. J Pharmacol Exp Ther 279(1), 69–76 doi:

Mattox, A.J., Carroll, M.E., 1996. Smoked heroin self-administration in rhesus monkeys. Psychopharmacology (Berl) 125(3), 195–201 doi:

Mattox, A.J., Thompson, S.S., Carroll, M.E., 1997. Smoked heroin and cocaine base (speedball) combinations in rhesus monkeys. Exp Clin Psychopharmacol 5(2), 113–118 doi:

McConnell, S.A., Brandner, A.J., Blank, B.A., Kearns, D.N., Koob, G.F., Vendruscolo, L.F., Tunstall, B.J., 2021. Demand for fentanyl becomes inelastic following extended access to fentanyl vapor self-administration. Neuropharmacology 182, 108355 doi: 10.1016/j.neuropharm.2020.108355.

Montanari, C., Kelley, L.K., Kerr, T.M., Cole, M., Gilpin, N.W., 2020. Nicotine e-cigarette vapor inhalation effects on nicotine & cotinine plasma levels and somatic withdrawal signs in adult male Wistar rats. Psychopharmacology (Berl) 237(3), 613–625 doi: 10.1007/s00213-019-05400-2.

Moore, C.F., Davis, C.M., Harvey, E.L., Taffe, M.A., Weerts, E.M., 2020. Antinociceptive, hypothermic, and appetitive effects of vaped and injected Δ9-tetrahydrocannabinol (THC) in rats: exposure and dose-effect comparisons by strain and sex. bioRxiv, 2020.2010.2006.327312 doi: 10.1101/2020.10.06.327312.

Morgan, D., Mitzelfelt, J.D., Koerper, L.M., Carter, C.S., 2012. Effects of morphine on thermal sensitivity in adult and aged rats. J Gerontol A Biol Sci Med Sci 67(7), 705–713 doi: 10.1093/gerona/glr210.

Moussawi, K., Ortiz, M.M., Gantz, S.C., Tunstall, B.J., Marchette, R.C.N., Bonci, A., Koob, G.F., Vendruscolo, L.F., 2020. Fentanyl vapor self-administration model in mice to study opioid addiction. Sci Adv 6(32), eabc0413 doi: 10.1126/sciadv.abc0413.

Nguyen, J.D., Aarde, S.M., Cole, M., Vandewater, S.A., Grant, Y., Taffe, M.A., 2016a. Locomotor Stimulant and Rewarding Effects of Inhaling Methamphetamine, MDPV, and Mephedrone via Electronic Cigarette-Type Technology. Neuropsychopharmacology 41(11), 2759–2771 doi: 10.1038/npp.2016.88.

Nguyen, J.D., Aarde, S.M., Vandewater, S.A., Grant, Y., Stouffer, D.G., Parsons, L.H., Cole, M., Taffe, M.A., 2016b. Inhaled delivery of Delta(9)-tetrahydrocannabinol (THC) to rats by e-cigarette vapor technology. Neuropharmacology 109, 112–120 doi: 10.1016/j.neuropharm.2016.05.021.

Nguyen, J.D., Grant, Y., Creehan, K.M., Hwang, C.S., Vandewater, S.A., Janda, K.D., Cole, M., Taffe, M.A., 2019. Delta(9)-tetrahydrocannabinol attenuates oxycodone self-administration under extended access conditions. Neuropharmacology 151, 127–135 doi: 10.1016/j.neuropharm.2019.04.010.

Nguyen, J.D., Grant, Y., Kerr, T.M., Gutierrez, A., Cole, M., Taffe, M.A., 2018a. Tolerance to hypothermic and antinoceptive effects of 9-tetrahydrocannabinol (THC) vapor inhalation in rats. Pharmacol Biochem Behav 172, 33–38 doi: 10.1016/j.pbb.2018.07.007.

Nguyen, J.D., Hwang, C.S., Grant, Y., Janda, K.D., Taffe, M.A., 2018b. Prophylactic vaccination protects against the development of oxycodone self-administration. Neuropharmacology 138, 292–303 doi: 10.1016/j.neuropharm.2018.06.026.

Pizzey, R., Hunt, N., 2008. Distributing foil from needle and syringe programmes (NSPs) to promote transitions from heroin injecting to chasing: an evaluation. Harm Reduct J 5, 24 doi: 10.1186/1477-7517-5-24.

Rook, E.J., Huitema, A.D., van den Brink, W., van Ree, J.M., Beijnen, J.H., 2006a. Pharmacokinetics and pharmacokinetic variability of heroin and its metabolites: review of the literature. Current clinical pharmacology 1(1), 109–118 doi:

Rook, E.J., van Ree, J.M., van den Brink, W., Hillebrand, M.J., Huitema, A.D., Hendriks, V.M., Beijnen, J.H., 2006b. Pharmacokinetics and pharmacodynamics of high doses of pharmaceutically prepared heroin, by intravenous or by inhalation route in opioid-dependent patients. Basic & clinical pharmacology & toxicology 98(1), 86–96 doi: 10.1111/j.1742-7843.2006.pto_233.x.

Smith, L.C., Kallupi, M., Tieu, L., Shankar, K., Jaquish, A., Barr, J., Su, Y., Velarde, N., Sedighim, S., Carrette, L.L.G., Klodnicki, M., Sun, X., de Guglielmo, G., George, O., 2020. Validation of a nicotine vapor self-administration model in rats with relevance to electronic cigarette use. Neuropsychopharmacology 45(11), 1909–1919 doi: 10.1038/s41386-020-0734-8.

Stephanie V. Ng, M.D., 2016. Opium Use in 19th-Century Britain: The Roots of Moralism in Shaping Drug Legislation. American Journal of Psychiatry Residents’ Journal 11(6), 14–14 doi: 10.1176/appi.ajp-rj.2016.110606.

Stover, H.J., Schaffer, D., 2014. SMOKE IT! Promoting a change of opiate consumption pattern -from injecting to inhaling. Harm Reduct J 11, 18 doi: 10.1186/1477-7517-11-18.

Strang, J., Griffiths, P., Gossop, M., 1997. Heroin smoking by ‘chasing the dragon’: origins and history. Addiction 92(6), 673–683; discussion 685-695 doi:

Vazda, A., Xia, W., Engqvist, H., 2021. The use of heat to deliver fentanyl via pulmonary drug delivery. Int J Pharm X 3, 100096 doi: 10.1016/j.ijpx.2021.100096.

Vendruscolo, J.C.M., Tunstall, B.J., Carmack, S.A., Schmeichel, B.E., Lowery-Gionta, E.G., Cole, M., George, O., Vandewater, S.A., Taffe, M.A., Koob, G.F., Vendruscolo, L.F., 2017. Compulsive-like sufentanil vapor self-administration in the rat. Neuropsychopharmacology in press doi:

Vendruscolo, J.C.M., Tunstall, B.J., Carmack, S.A., Schmeichel, B.E., Lowery-Gionta, E.G., Cole, M., George, O., Vandewater, S.A., Taffe, M.A., Koob, G.F., Vendruscolo, L.F., 2018. Compulsive-Like Sufentanil Vapor Self-Administration in Rats. Neuropsychopharmacology 43(4), 801–809 doi: 10.1038/npp.2017.172.

Wang, Q.L., Liu, Z.M., 2012. Characteristics of psychopathology and the relationship between routes of drug administration and psychiatric symptoms in heroin addicts. Subst Abus 33(2), 130–137 doi: 10.1080/08897077.2011.630945.

